# Deep profiling reveals substantial heterogeneity of integration outcomes in CRISPR knock-in experiments

**DOI:** 10.1101/841098

**Authors:** Hera Canaj, Jeffrey A. Hussmann, Han Li, Kyle A. Beckman, Leeanne Goodrich, Nathan H. Cho, Yucheng J. Li, Daniel A. Santos, Aaron McGeever, Edna M. Stewart, Veronica Pessino, Mohammad A. Mandegar, Cindy Huang, Li Gan, Barbara Panning, Bo Huang, Jonathan S. Weissman, Manuel D. Leonetti

## Abstract

CRISPR/Cas technologies have transformed our ability to add functionality to the genome by knock-in of payload via homology-directed repair (HDR). However, a systematic and quantitative profiling of the knock-in integration landscape is still lacking. Here, we present a framework based on long-read sequencing and an integrated computational pipeline (knock-knock) to analyze knock-in repair outcomes across a wide range of experimental parameters. Our data uncover complex repair profiles, with perfect HDR often accounting for a minority of payload integration events, and reveal markedly distinct mis-integration patterns between cell-types or forms of HDR templates used. Our analysis demonstrates that the two sides of a given double-strand break can be repaired by separate pathways and identifies a major role for sequence micro-homology in driving donor mis-integration. Altogether, our comprehensive framework paves the way for investigating repair mechanisms, monitoring accuracy, and optimizing the precision of genome engineering.

## Introduction

Recent developments in gene editing technologies have transformed our ability to manipulate genomes. Programmable site-specific nucleases – in particular CRISPR/Cas systems – introduce double-strand breaks (DSBs) at chosen genomic locations, prompting the activation of two separate DNA repair pathways which can be leveraged for genome engineering^1^. On the one hand, non-homologous end-joining (NHEJ) can introduce insertions or deletions (in-dels) at the DSB site to inactivate gene products or regulatory elements. On the other hand, homology-directed repair (HDR) can use exogenous DNA sequences as templates to integrate (knock-in) new genetic information in the locus of interest^2^. Knock-in strategies have wide applications ranging from correcting disease-causing mutations or inserting therapeutic payloads in a clinical context^3–5^, to introducing functional reporters for cell biology research^6, 7^.

Ultimately, the power of genetic engineering will rely on our ability to predictably edit genomes in order to precisely control cellular behaviors. Traditionally, NHEJ and HDR have been set apart by their degree of predictability: NHEJ is often thought to drive random repair outcomes, while HDR is considered to enable precise and templated editing. Recent data is challenging this simple distinction, however. High-throughput sequencing of NHEJ repair has uncovered complex but actionable sequence patterns that control in-del outcomes at DSBs^8–13^. In some cases, predictable NHEJ outcomes can be leveraged to precisely correct pathogenic human mutations, paving the way for template-free therapeutic genome editing^9^. By contrast, while accumulating evidence suggests that the integration of payload in knock-in experiments is not always precise^7, 14, 15^, a systematic and quantitative profiling of the full spectrum of HDR repair outcomes is still missing.

Given its wide range of applications in both clinical and research settings, understanding the parameters that govern the efficiency and precision of knock-in editing is critical. To what extent is HDR precisely templated? How are the profiles of knock-in outcomes influenced by target loci, payload size or the form of DNA donor template being used (e.g., single- or double-stranded)? Are these parameters equally impactful across different cell types? Here, we present a comprehensive set of experimental workflows and computational tools for the deep profiling of knock-in repair in mammalian cells to address these salient questions. Our data uncovers a surprising degree of repair complexity, which is both donor type- and cell type-dependent, and reveals that different mechanisms often interplay for the repair of a given allele.

## Results

### *knock-knock*: a versatile tool for mapping repair architecture

We initially sought to build solutions for the high-throughput genotyping of loci integrated with fluorescent proteins for cell biology applications (Figure 1A). We reasoned that improvements in Single-Molecule Real-Time (SMRT, Pacific Biosciences) sequencing chemistry^16^ would allow for the rapid and accurate long-read sequencing of repaired alleles from amplicon libraries (Suppl. Figure 1A-C). We first compared the performance of PCR, ssDNA and plasmid donors for the integration of GFP at the RAB11A N-terminus in HEK293T cells (Figure 1B). After sorting polyclonal pools of fluorescent cells (GFP+) by flow cytometry to enrich for knock-in alleles, we generated amplicon libraries from cell lysates followed by SMRT sequencing and circular consensus sequence reconstruction. Surprisingly, the distribution of amplicon length in GFP+ cells revealed a diversity of sizes, many of which did not match the expected length for HDR or wild-type (wt) ± in-dels alleles (Figure 1B). To explore the complex landscape of editing outcomes implied by this diversity, we developed *knock-knock*, a computational framework for characterizing the allelic architectures produced by knock-in editing experiments. The central goal of *knock-knock* is to produce an unbiased taxonomy of editing outcomes from amplicon sequencing data, providing a more granular description of allele diversity than existing software tools allow^17, 18^. To accomplish this, the algorithm makes minimal assumptions about the possible configurations of sequence in each amplicon. Instead, for each sequencing read, *knock-knock* generates a comprehensive set of local alignments between segments of the read and all possible sequences present in the experiment: the targeted genome, the full HDR donor template, and any other relevant sequences that are provided. It then identifies a parsimonious subset of local alignments that covers the amplicon and parses the arrangement of these alignments to classify the editing outcome (Figure 1C). To facilitate interpretation of these outcomes, *knock-knock* produces interactive visualizations for exploring the read architectures identified in an experiment. *knock-knock* supports data from both long read (SMRT) and short read (paired-end Illumina) sequencing platforms, and is available as an open-source software package at https://github.com/jeffhussmann/knock-knock.

**Figure 1:**
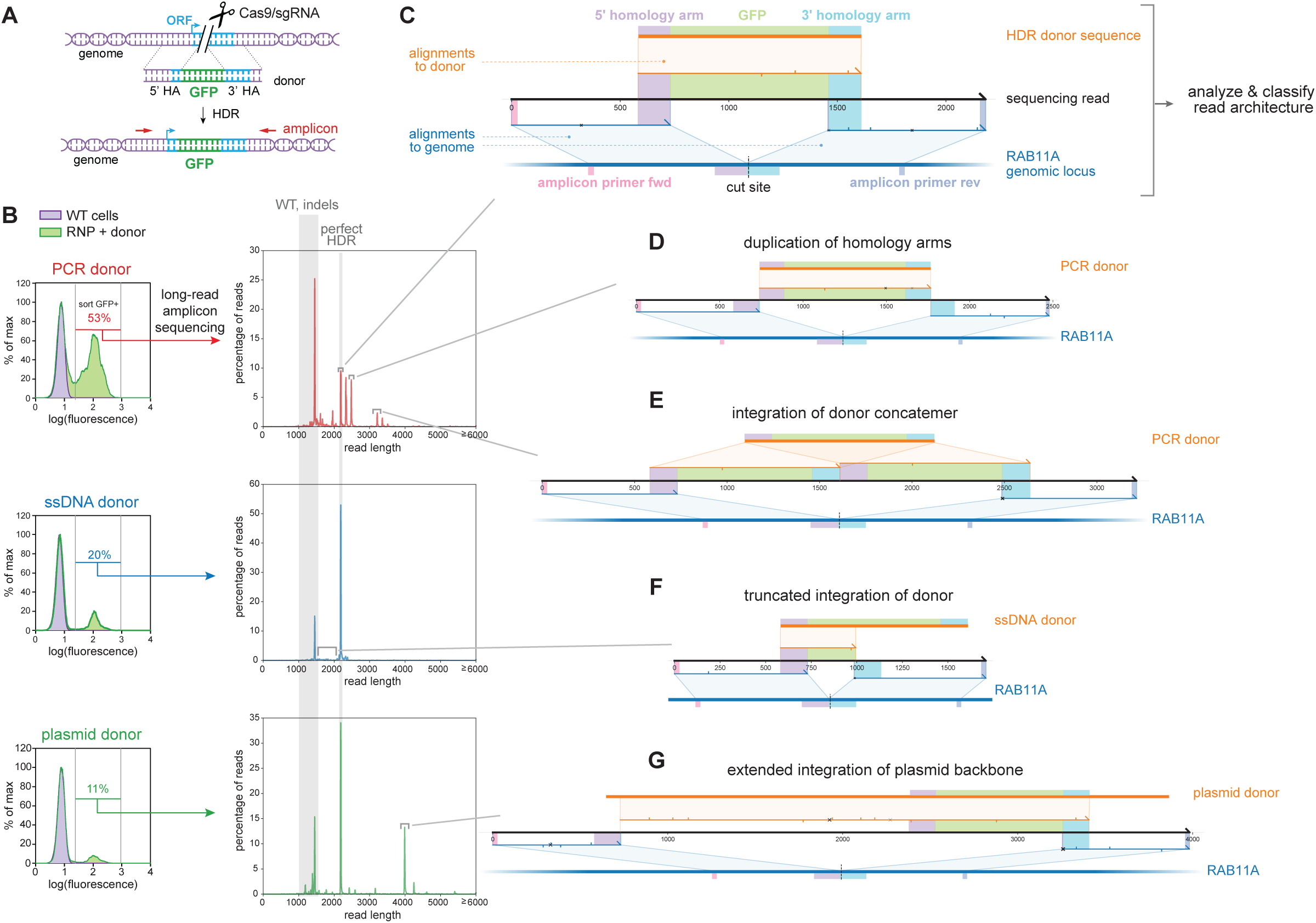
knock-knock analysis of repair architecture. **A**. Experimental design: HDR knock-in of GFP in open-reading frames (ORFs), followed by amplicon sequencing. **B**. (Left) Flow-cytometry fluorescence profiles of GFP insertion at the RAB11A N-terminus in HEK293T cells using plasmid, PCR or long ssDNA donors. (Right) Corresponding distribution frequency of amplicon length after selection of GFP+ cells and SMRT long-read amplicon sequencing of RAB11A. Lengths corresponding to wt ± in-dels and expected HDR product are shaded in grey. **C**. Example of *knock-knock* classification. Each sequencing read (black) is deconstructed through a series of local alignments to either the reference genome (blue) or any other sequence provided, in particular the donor template (orange). The architecture of each read is further analyzed and classified into phenotypic outcomes. Final results are available within an interactive visualization framework including relevant sequence boundaries (forward and reverse sequencing primers, homology arms and GFP payload) are highlighted in colors. Sequence alignment mismatches between sequencing read and reference sequences are indicated: cross for sequence mismatch, down-tick for 1-bp insertions, up-tick for deletions. **D-E**. Examples of mis-integration identified from cells engineered using PCR donor: blunt insertion resulting in duplication of the homology arms, and integration of donor concatemer. **F**. Examples of mis-integration identified from cells engineered using ssDNA donor: truncated insertion of GFP payload. **G**. Example of mis-integration identified from cells engineered using plasmid donor: integration of donor plasmid backbone sequence.

Applying *knock-knock* to SMRT amplicons from the RAB11A-GFP+ cells revealed a striking diversity of mis-integration patterns that differed across donor types (Figure 1D-F). First, cells edited using double-stranded PCR donors exhibited a mis-integration profile dominated by ligation of the donor ends into the genomic cut site or onto one another, leading to alleles with duplicated homology arms (Figures 1D and S1E) or with integration of multiple concatenated copies of insert sequences (Figures 1E and S1F). Ligation of reactive dsDNA ends into a genomic DSB is an expected phenomenon which has been leveraged by methods for profiling off-target sites of Cas9 gRNA cleavage^19^, and can lead to concatemerization when donor fragments react with one another^20, 21^. Second, cells edited using long single-stranded donors (produced by reverse-transcription of an RNA intermediate using a thermo-stable group II reverse transcriptase^22^, see Material and Methods) showed the integration of truncated GFP payload sequences (Figure 1F). Finally, cells edited using plasmid HDR donor displayed a strong signature of integration of plasmid backbone sequence past homology arm boundaries (Figures 1G and S1H). Interestingly, high rates of backbone integration from plasmid donors have been previously reported in a systematic study of gene editing in human induced pluripotent stem cells (iPSCs)^7^, and in the genome of cattle engineered using plasmid HDR templates^23, 24^. Of note, *knock-knock* can also quantify a variety of additional editing outcomes including spurious integration of exogenous DNA sequences (such as rare integrations of *E. coli* genomic DNA presumably carried over from plasmid preparation, Figure S1G) or longer range deletions^25^ which often arise at micro-homology overlaps (Figure S1I). Altogether our initial analysis – enabled by *knock-knock* – highlighted a broad complexity of mis-integration profiles in CRISPR knock-in experiments, with different signatures between donor types.

### Deep profiling of repair profiles across donor- and cell-types

Our initial results motivated a more systematic characterization of integration outcomes across a wide range of experimental conditions (size of insert, locus, donor-type and cell-type). We chose electroporation of pre-complexed *S. pyogenes* Cas9/gRNA complexes as a delivery system given its prevalence, breadth of implementation and lower off-target activity^26, 27^. To encompass a maximal diversity of donor types, we sought to profile integration of both small (<100 nt) and large (∼1 kb) payloads because payload size typically informs the type of donor used as template for HDR: small payloads usually involve synthetic single-stranded oligonucleotide donors (ssODNs, commercially available under 200nt) while gene-sized integrations often use plasmid or PCR-generated double-stranded donors (dsDNA) ^5, 7, 14, 28^. To take advantage of fluorescent readouts to monitor HDR efficiency, we leveraged the GFP11 split-GFP system^6^ for small payload integration (90nt payload, 55 nt homology arms), while using the full-length super folder GFP (sfGFP) for gene-sized insertions (∼700nt payload, ∼150-600nt homology arms which we show supply sufficient homology to drive efficient knock-in, Suppl. Figure 2A). Finally, because DNA repair is known to vary between cell types^29^, we sought to compare integration profiles in both transformed (HEK293T) and non-transformed (human iPSCs, mouse embryonic stem cells – mESCs) cell lines. In HEK293T, we took advantage of cell-cycle synchronization with nocodazole treatment to promote HDR^26^.

We first examined GFP11 integration in HEK293T and iPSCs with ssODN, PCR and plasmid donors, measuring integration efficiency by fluorescence flow cytometry. Across conditions, the amount of donor deliverable into cells is inherently limited by synthesis yield (eg. PCR or plasmid preparation) and toxicity with the exception of synthetic ssODNs, which are readily available in large amounts and exhibit very low toxicity^5^. At equimolar amounts, we found that PCR donors enabled highest GFP11 integration in 4 loci tested (Figure 2A). ssODN performance could be significantly improved by increasing the quantity of donor delivered, making ssODN the most efficient form of donors for small insertions (Figure 2A). Because single-stranded and PCR donors showed the best performance overall, we further compared their behavior for integration of the large sfGFP payload. While both donor types could drive efficient integration, PCR donors consistently displayed the highest efficiency across cell-types and loci (Figure 2B).

**Figure 2:**
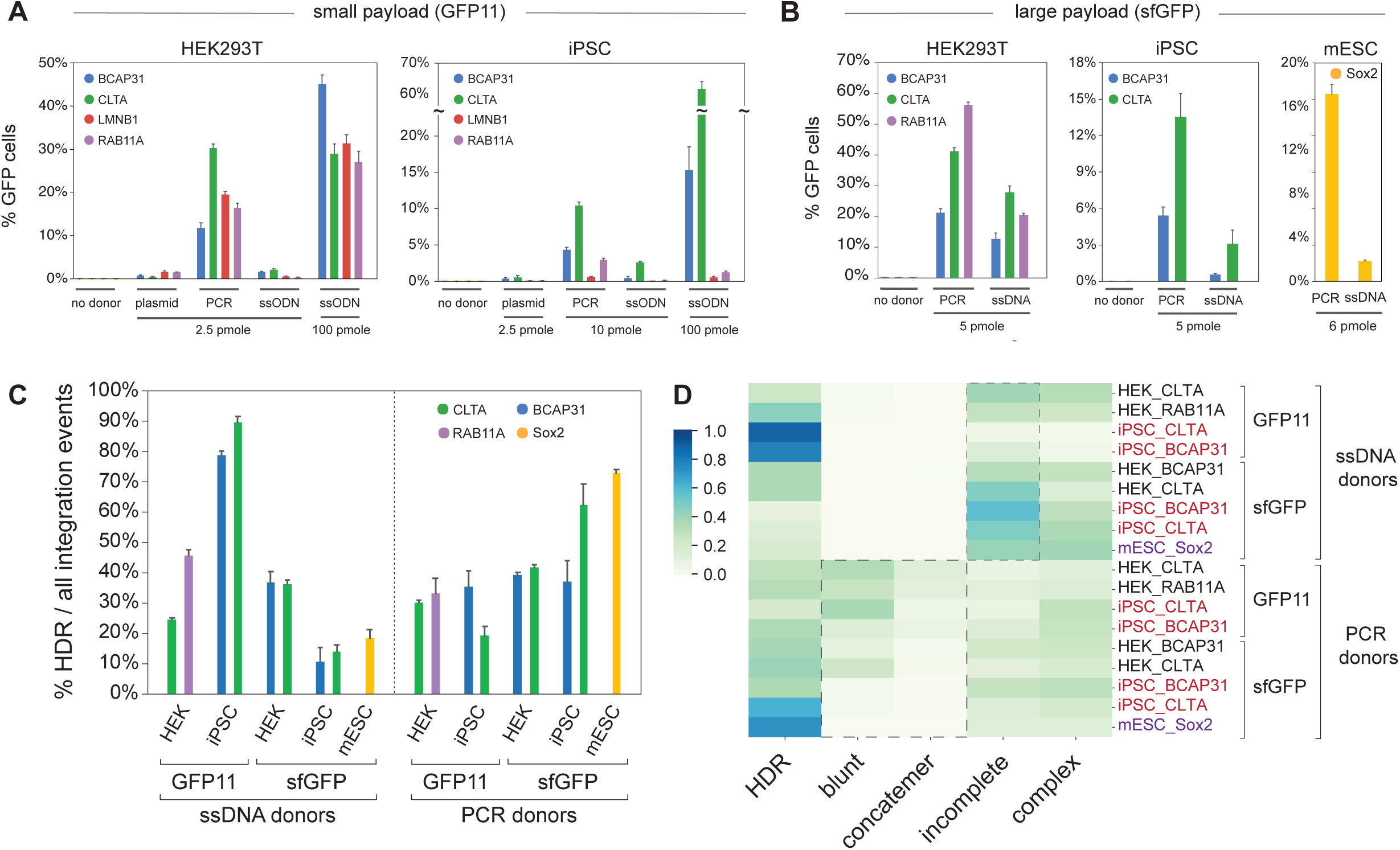
Deep profiling of integration signatures. **A.** Functional read-out of knock-in efficiency of small GFP11 payload (90 nt). Percentage of fluorescent cells edited using different donor types (plasmid, PCR, ssODN) is quantified by flow cytometry in HEK293T and iPSC cells. The amount of donor electroporated for each 200,000 cells is specified. Shown are mean + standard deviation from 3 independent replicate experiments (some error bars are too small to be visible). **B.** Functional read-out of knock-in efficiency of large sfGFP payload (∼700 nt). Percentage of fluorescent cells edited using different donor types (PCR, ssDNA) is quantified by flow cytometry in HEK293T, iPSC and mESC cells. The amount of donor electroporated for each 200,000 cells is specified. Shown are mean + standard deviation from 3 independent replicate experiments (some error bars are too small to be visible). **C.** Prevalence of HDR within all integration events (repair events in which donor sequences were found integrated in the target locus). Shown are mean + standard deviation from 3 independent replicate experiments. **D.** Heatmap representing the distribution of integration events, classified across different repair categories. Each line represents average values from one experimental condition. Related to Suppl. Fig. 2C-D.

On the basis of these efficiency readouts, we focused on single-stranded and PCR donors for further profiling of integration outcomes by SMRT amplicon deep sequencing and *knock-knock* analysis. To facilitate phenotypic description, we classified these outcomes into 5 functional categories (Suppl. Figure 2B): “HDR” alleles, the expected HDR products; “blunt” alleles, which contain the direct ligation of donor ends on at least one side of the genomic DSB; “concatemer” alleles, which show insertion of multiple donor sequences ligated together; “incomplete” alleles, which display a correct HDR junction for only one of the two homology arms (excluding any “blunt” ligations on the other arm, which are classified above); and “complex” alleles, which do not fall into any of the previous categories. To capture the complete diversity of integration events, we profiled genotypes from the entire gene-edited pool without any selection (i.e., without fluorescent cell sorting). In HEK293T and iPSC we profiled two loci for each donor type.

Three main conclusions stand out from the detailed analysis of all experimental conditions, which is shown in Suppl. Figures 2C-D. First, the landscape of integration in CRISPR knock-in experiments is complex: perfect HDR represents only a minority of integration events in most cases (Figure 2C) and different mis-integration profiles are observed between conditions (Figure 2D). ssODN donors in iPSCs (small payload integration) are an exception, with >75% of integration events being HDR (Figure 2C). Highlighting the complexity we uncovered, ssDNA donors performed conversely very poorly for large payload integration in iPSCs (<20% perfect HDR, Figure 2C), revealing that payload size might affect integration outcomes. Second, cell types can behave differently within the same set of conditions. For example, the high HDR specificity of ssODN-mediated integration seen in iPSC is not observed in HEK293T, even for the same gene (CLTA). Third, while clear patterns are hard to delineate from the heterogeneity of the profiles, two signatures differentiate ssDNA from PCR donors: PCR donors have a propensity to cause blunt or concatemeric integrations, while the profiles from ssDNA donors are dominated by incomplete integrations (Figure 2D). Note that in HEK293T we verified that the complexity integration profiles we observe did not originate from cell-cycle synchronization (Suppl. Figure 2E). Overall, our systematic analysis reveals a broad diversity of repair outcomes in CRISPR knock-in experiments and the association of distinct repair profiles with cell type and donor class.

### Mechanistic insights into non-HDR integrations

Next, we sought to understand the underlying mechanisms driving each particular class of non-HDR events (subsequently referred to as “mis-integration”). With PCR donors, one mechanism is apparent from the high rate of blunt and concatemer integrations we observed: an NHEJ-like ligation of donor molecules into genomics DSBs (or ligation of donors onto themselves, for concatemer reads). Interestingly, this is a major mis-integration mechanism for small payload in both HEK293T and iPSC, and for large payload in HEK293T. By contrast, this is not very prevalent for large payload mis-integration in iPSC or mESC (Suppl. Figure 2D), highlighting again differences in DNA repair profiles between cell lines and donor types. From such NHEJ-based mechanisms, a testable hypothesis is that modifications of the donor 5’ ends to protect reactive hydroxyl groups should mitigate mis-integration. Focusing on the small GFP11 payload, we designed two forms of 5’ modified PCR donors (Figure 3A): a version with bulky biotin molecules, and a “hybrid” design containing artificial single-stranded ends isolated by a PEG linker (seeking to take advantage of the fact that ssDNA donors give rise to negligible levels of blunt ligations, see Suppl. Figure 2C). As expected, 5’ modifications significantly decreased ligation-based mis-integration in both HEK293T and iPSC (Figure 3B) and increased the relative levels of correct HDR compared to total integration events (Suppl. Figure 3). Altogether, 5’ end modifications are a useful strategy to decrease ligation-based mis-integration of PCR donors (which has been leveraged previously for genome editing in medaka fish^20^) but fail to completely resolve mis-integration events.

**Figure 3:**
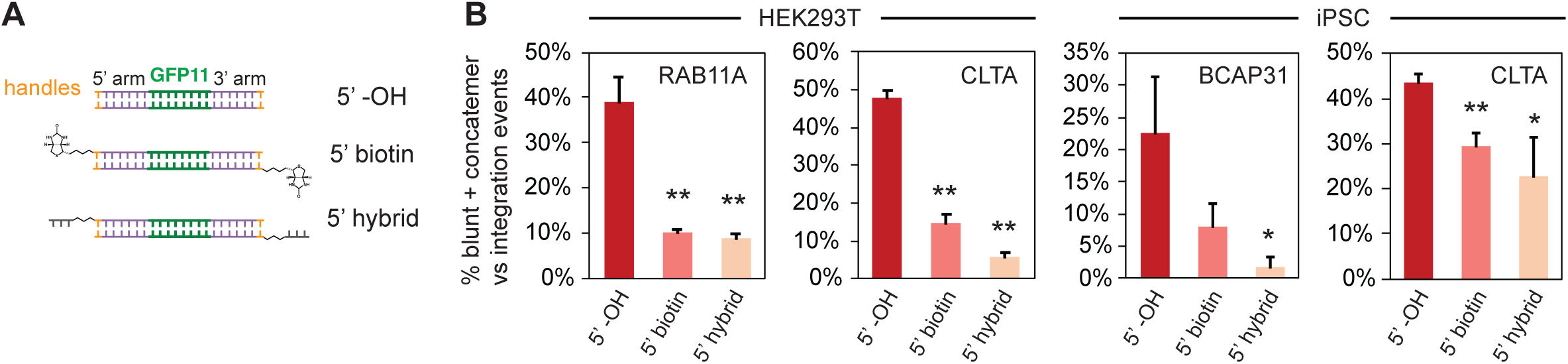
5’ modifications mitigate mis-integration from PCR donors. **A.** Modifications used for GFP11 donors: 5’ biotin and 5’ hybrid design in which single-stranded DNA ends are introduced after a PEG spacer. **B.** Prevalence of blunt + concatemeric reads within all integration events (GFP11 integration in HEK293T and iPSC). Shown are average + standard deviation from 3 independent replicate experiments. 5’ modifications alleviate blunt/concatemeric mis-integrations across experimental conditions. * p≤0.05, ** p≤0.01, unpaired Student’s t-test compared to 5’ -OH control.

Another prevalent hallmark of mis-integration is that a large number of these events display a proper HDR junction at one of the two homology arms, while repair at the other homology arm resolves through a non-HDR mechanism (“asymmetric HDR”, Figure 4A and Suppl. Figure 4A). Such asymmetric HDR has been observed before^14, 15, 30^ but has never been quantified across a wide range of conditions. Asymmetric HDR tends to be more pronounced with ssDNA donors (50-80% of total integration events depending on specific conditions used) than with PCR donors (25-50%) (Figure 4A, Suppl. Fig. 4A). With ssDNA donors, HDR asymmetry displays a strong directionality with respect to the 5’ to 3’ direction of the donor itself. We defined a directionality factor as: [fraction of asymmetric HDR reads that have a non-HDR junction oriented on the 3’ side of the donor] - [fraction of asymmetric HDR reads that have a non-HDR junction oriented on the 5’ side of the donor]. For ssDNA donors, asymmetric HDR was dominated by reads showing imprecise junction on the 5’ side (directionality around +1.00, Figure 4B). By comparison, asymmetric HDR did not have a specific directionality for PCR donors overall (Figure 4B, “direction” of PCR donors was defined based on the corresponding ssDNA conditions). The directionality we observe for ssDNA donors can be understood in the context of the synthesis-dependent strand-annealing model that has been proposed for ssDNA-mediated repair^14^, where the 3’ end of ssDNA donors must first anneal to single-stranded genomic sequences that are left exposed after DSB resection (Figure 4C). This results in a correct HDR junction at the 3’ side of the donor, while repair can be resolved in a non-HDR fashion at the other end of the DSB (Figure 4C). With PCR donors, both donor strands can act independently and have an equal chance of engaging DNA repair, leading to an overall loss of directionality.

**Figure 4:**
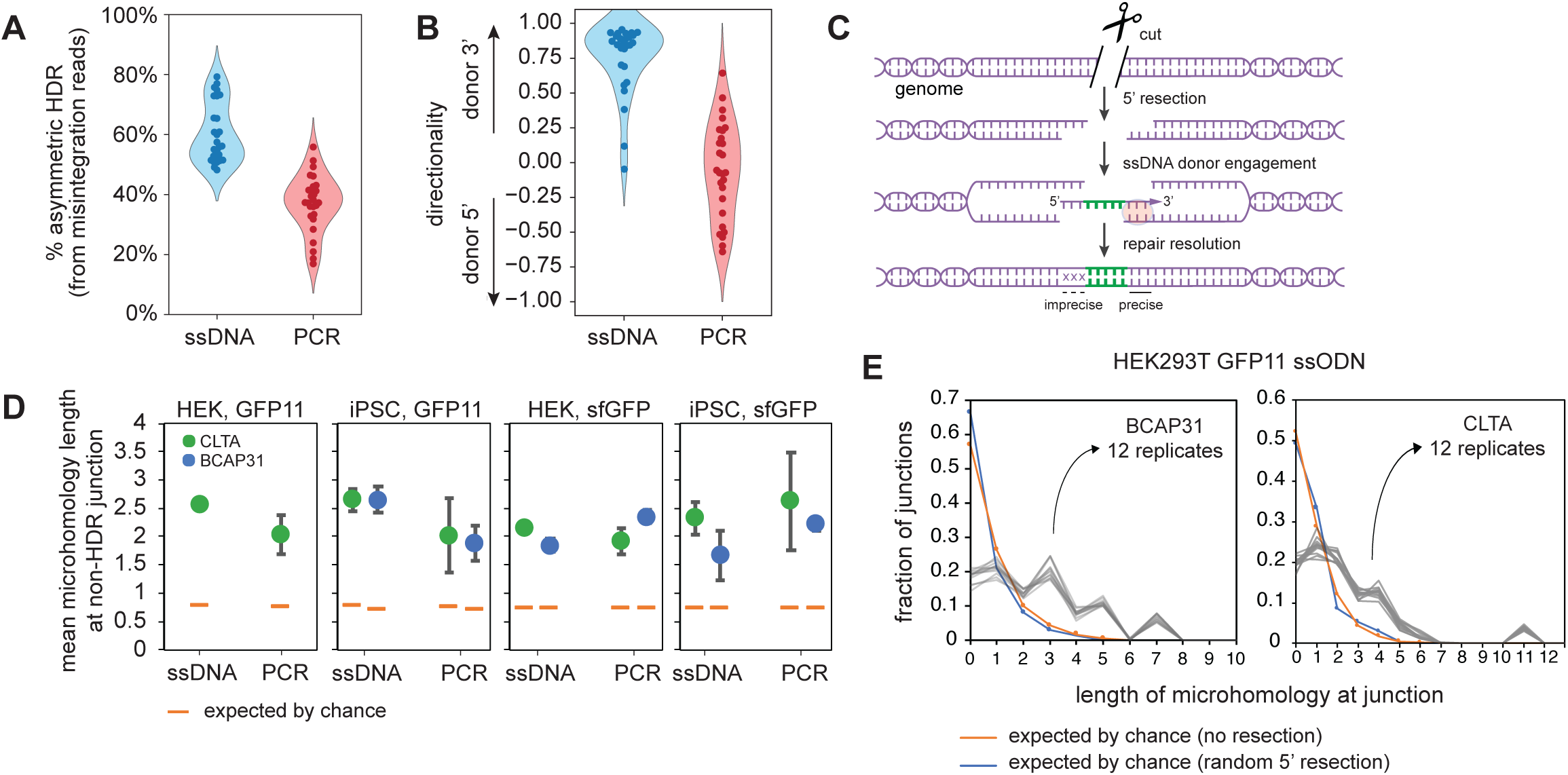
asymmetric HDR and microhomology-mediated repair. **A.** Prevalence of asymmetric HDR (defined as repair events showing a proper HDR junction at only one homology arm) in mis-integration reads across experimental conditions. Related to Suppl. Fig 4A. **B.** Directionality of asymmetric repair from incomplete reads. **C.** Synthesis-dependent strand-annealing model of ssDNA-mediated DSB repair. **D.** Quantification of the mean length of microhomology between payload and genome sequences observed at non-HDR junctions from incomplete reads (SMRT sequencing). Shown are average and standard deviation from 3 independent replicate experiments. The average microhomology length expected by chance is shown as an orange bar. Note: BCAP31 GFP11 integration was not measured in HEK293T. **E.** Distribution of microhomology length at incomplete non-HDR junctions across 12 replicate experiments (BCAP31 and CLTA loci, ssODN GFP11 integration in HEK293T, MiSeq sequencing). The distributions of microhomology expected by chance considering non-resected or randomly resected DSBs are shown in orange and blue, respectively.

**Figure 5:**
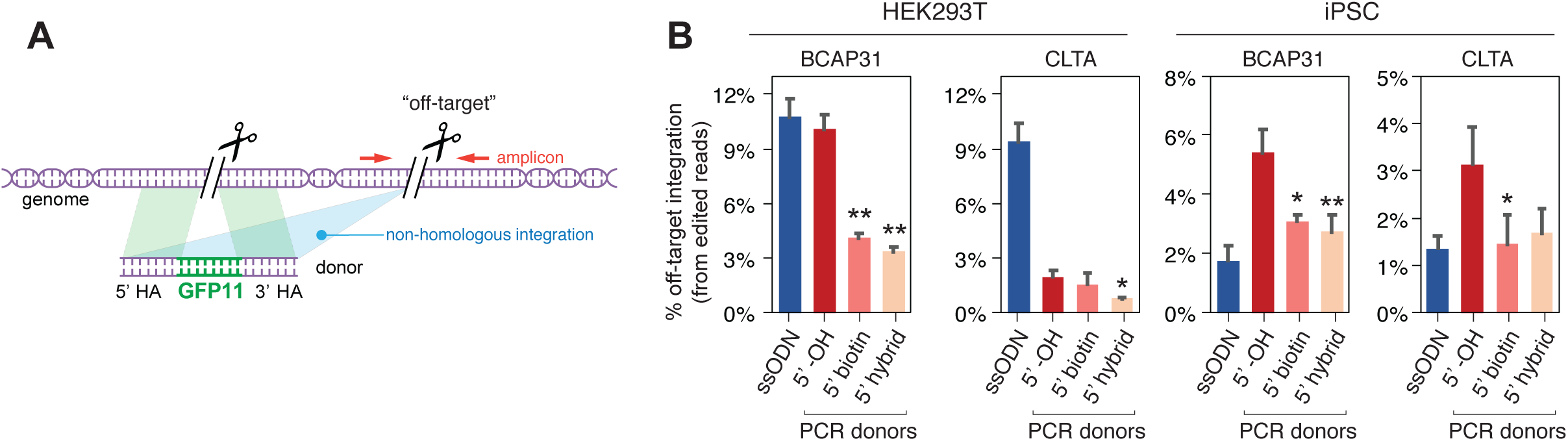
off-target integration across donor types. **A.** Experimental design. Genome is cut at two separate loci while a GFP11 homology donor is provided for only one locus. Any integration of the donor at the non-homologous cut site represents an “off-target” integration, which is quantified through amplicon sequencing at the off-target site. **B.** Quantification of off-target integration at each specified locus, for ssODN and PCR donors in both HEK293T and iPSC. Integration is normalized to edited (non-wt) reads. 5’ end modifications of PCR donors alleviate off-target integration. Shown are average + standard deviation from 3 independent replicate experiments. * p≤0.05, ** p≤0.01, unpaired Student’s t-test compared to 5’ -OH control.

Given the prevalence of HDR asymmetry, we sought to characterize the mechanisms by which repair is resolved at the “non-HDR” homology arm junction. Because direct 5’ end ligation could only explain a fraction of such asymmetric events (Suppl. Figure 4B), we further examined other DNA repair mechanisms. Strikingly, we discovered a major role for microhomology in driving non-HDR junctions in asymmetric reads. Across conditions, such junctions often occurred at sequence locations where donors and genome shared microhomology (Suppl. Figure 4C) and the length of microhomology at observed junctions was significantly higher than expected by chance across experimental conditions (Figure 4D). We reasoned that since microhomology is solely dictated by sequence details at a given locus, patterns of microhomology-driven mis-integration should be very reproducible within a given experimental condition (locus, payload, cell type). To test this, we analyzed mis-integration at two separate loci over 12 independent replicate experiments using GFP11 payload with ssODN donors in HEK293T cells. In this experimental setup, asymmetric mis-integration events can be fully captured within <600 bp amplicons which allow the use of MiSeq Illumina sequencing for profiling read architecture with higher fidelity and throughput than SMRT sequencing. This analysis revealed a remarkable reproducibility of mis-integration profiles across replicates (Figure 4E), confirming the prevalence of sequence microhomology in dictating asymmetric repair. Similar results were obtained in iPSC (Suppl. Figure 4D). Finally, we verified that these integration patterns did not originate from truncated synthetic products of the ssODNs we used as donors. Indeed, gel-purified ssODN donors revealed similar repair profiles (Suppl. Figures 4E-G), confirming that microhomology (and not artificial truncation of donors) is a major driving force behind incomplete integration.

### Non-homologous integration of donors at off-target cut sites

Finally, we analyzed an undesirable by-product of knock-in: the possible integration of donor sequences at unintended locations in the genome, typically at off-target nuclease cutting sites^15, 31^. For this, we compared the propensity of single-stranded and PCR donors to integrate in a non-homologous fashion by designing experiments where the genome is cut concomitantly at two separate positions (ie. using two targeting gRNAs). Co-delivery of donor containing homology to only one of the two cut sites creates a situation where the second site acts as an artificial “off-target” cut, allowing quantification of non-homologous integration by SMRT sequencing and *knock-knock* (Figure 4A). To correct for different Cas9 cutting efficiencies between conditions, off-target integrations were normalized for total editing at the corresponding site (ie. non-wt reads). We monitored integration of a GFP11 payload in HEK293T and iPSC at two different “off-target” loci (CLTA and BCAP31), and found that the propensity of non-homologous integration varied between the two cell types (Figure 4B). In particular, ssODN donors showed high level of non-homologous integration in HEK293T (∼10%) but not in iPSC (∼1.5%), mirroring the difference we observed for on-target integration. PCR donors showed more variable integration rates between the two loci tested. Interestingly, 5’ modifications could significantly decrease off-target integration of PCR donors in both cell types, showing that 5’ end capping might have a general benefit beyond promoting higher HDR rates.

## Discussion

We present *knock-knock*, an open-source software package for characterizing and quantifying the full diversity of repair outcomes in CRISPR-based knock-in experiments. Applying *knock-knock* to knock-in experiments performed across a broad range of experimental conditions uncovers a multi-faceted complexity of DNA repair outcomes. Strikingly, we observe that bona fide HDR represents only a fraction of integration events in the conditions we tested, with pronounced differences between cell types and donor types (Figure 2D). Interestingly, a majority of mis-integration events still contain a proper HDR junction at one of the two homology arms (Figure 4A), highlighting that both homologous and non-homologous mechanisms can compete for repair of the same allele^14, 30^. In line with previous studies^20, 21^, we show that NHEJ-based ligation is a major by-product of knock-in with linear dsDNA donors, which can be alleviated by chemical modifications of the DNA ends. Importantly however, classical NHEJ is not the only repair mechanism competing with HDR for DSB resolution. Indeed, our results reveal a major contribution of microhomology-mediated repair in mis-integration events that is especially pronounced with ssDNA donors. This underlines the broad importance of microhomology patterns in shaping DNA repair, as recent studies also unveiled a major role for microhomology in shaping in-del profiles in knock-out experiments^8–13^. Notably, some knock-in strategies leverage NHEJ or microhomology to integrate payload with minimal (or absent) homology arms^32–34^ to overcome inherent limitations of HDR, in particular its requirement of active cell cycle progression. How precisely repair outcomes can be controlled in such strategies remains under-explored, but our profiling approach should facilitate their deeper characterization.

The main conclusion from our current results is that knock-in integration landscapes can be very heterogeneous and condition-dependent, and therefore need to be characterized more systematically. The recent debate surrounding the discovery of plasmid backbone sequences in the genome of engineered cattle^23, 24^ (derived from mis-integration of an HDR donor template) also highlights the universal need for comprehensive genotyping post-editing. Most existing analytical procedures seek to assess editing conditions based on a limited set of heuristic variables (e.g. % GFP+ cells, digital PCR). By contrast our approach (SMRT amplicon sequencing and *knock-knock* software) allow experimenters to relate editing conditions with genotypic outcomes, potentially allowing for the optimization towards specific genotypic profiles. While SMRT sequencing has long been recognized as a powerful genotyping approach^35^, recent advances in SMRT technology now enable both higher accuracy in consensus base calling (allowing for a dissection of repaired alleles at nucleotide resolution) and higher throughput^16^. Note that to profile small edits or insertions, Illumina amplicon sequencing (up to 600bp length on a paired-end MiSeq run) might also be appropriate – this is especially relevant for ssODN donors, where incomplete integrations dominate so that short amplicon lengths might still be able capture the whole diversity of integration events. For all applications, our open-source *knock-knock* package enables the precise profiling of DNA repair with great flexibility in experimental design.

Altogether, the strong dependence of knock-in profiles on cell- and donor-types calls for a much more exhaustive analysis of experimental conditions than the scope of a single study permits. From a practical standpoint, the choice of donor-type for a specific application will have to balance both efficiency and precision of integration. From our current data (Suppl. Figure 5), we think that ssODN donors are an attractive option for the integration of small payload (especially given that they require no preparation) while PCR donors (potentially using 5’ end modifications) might be advantageous for larger knock-ins. It would be very interesting to test whether HDR precision can be further optimized by forms of donors we did not test here. Viral-based donors are an attractive possibility. For example, recombinant adenovirus donors have been found to prevent concatemerization of knock-in integration on the basis of their 5’ covalent linkage with endogenous terminal proteins^21^; whether these can also prevent other forms of mis-integration remains to be tested. Finally, while we only profiled DSBs triggered by the classical *S. pyogenes* Cas9 system, a whole arsenal of designer nucleases is now available (including different Cas proteins, nickases, modified Cas and non-Cas systems), complemented recently by knock-in strategies that do not require DSBs^36–38^. We anticipate that the interactive exploration of editing outcomes enabled by *knock-knock* will be broadly applicable to understanding how these different approaches might shape different repair outcome profiles. Overall, our study highlights a complexity of competing pathways at play for the integration of payload in genome editing experiments and paves the way for a more complete elucidation of the underlying cellular mechanisms. It will be fascinating to see whether “surgical” adjustments of cellular factors (e.g. inhibiting or activating endogenous repair pathways) or of experimental conditions (e.g. specific donor types or nucleases) will enable the optimization of precisely templated knock-in integration for both research and clinical applications.

## Supporting information

Supplementary folder 1

## Acknowledgements

We thank the Genomics team at CZ Biohub (Norma Neff, Michelle Tan, Rene Sit) and Shana McDevitt (Vincent J. Coates Genomics Sequencing Laboratory, UC Berkeley) for help with deep-sequencing, and members of the Weissman and Leonetti labs for comments. M.D.L. thanks Chan Lek Tan for critical feedback. J.A.H. is the Rebecca Ridley Kry Fellow of the Damon Runyon Cancer Research Foundation (DRG-2262-16).

## FIGURE LEGEND – SUPPLEMENTARY

**Supplementary Figure 1 (related to Fig. 1):**
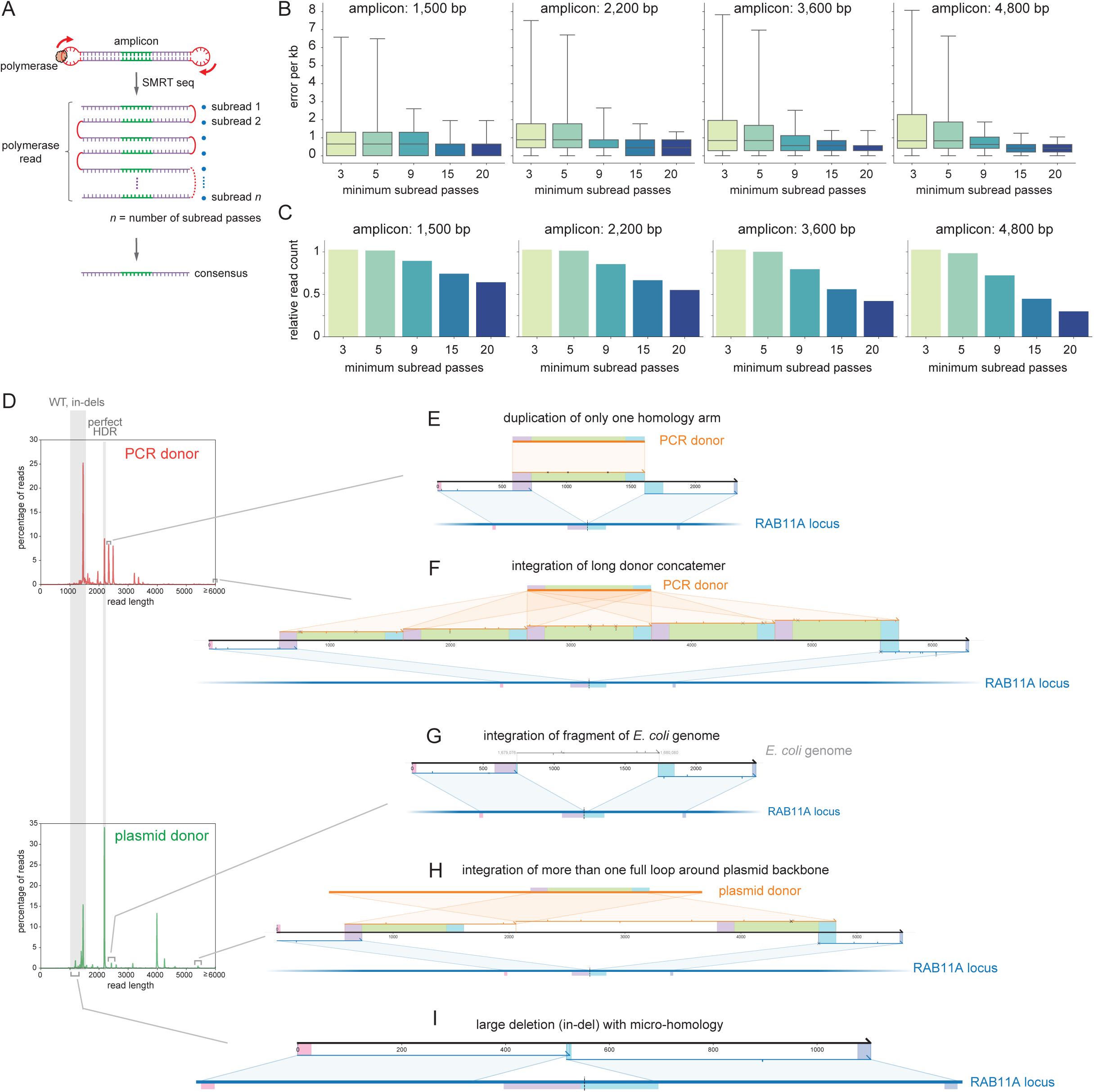
SMRT sequencing and *knock-knock* analysis. **A.** SMRT sequencing. A given amplicon is sequenced across many subreads, from which a consensus sequence can be computed. Individual polymerase reads can have varying numbers (n) of subreads passes. **B.** Measurement of SMRT sequencing error rate after consensus calling, using reference amplicons of different lengths (1,500-4,800 bp). Sequencing error rates per kb can be decreased by limiting consensus calling to reads with increasing minimum number of subread passes. Boxes represent median + 25th and 75th percentiles; whiskers represent 5th and 95th percentiles. **C.** Barplot showing how subread passes stringency impacts the number of polymerase reads considered. **D-I.** Related to Fig. 1B-G. Examples of reads identified from *knock-knock* analysis of HEK293T cells edited

**Supplementary Figure 2 (related to Fig. 2):**
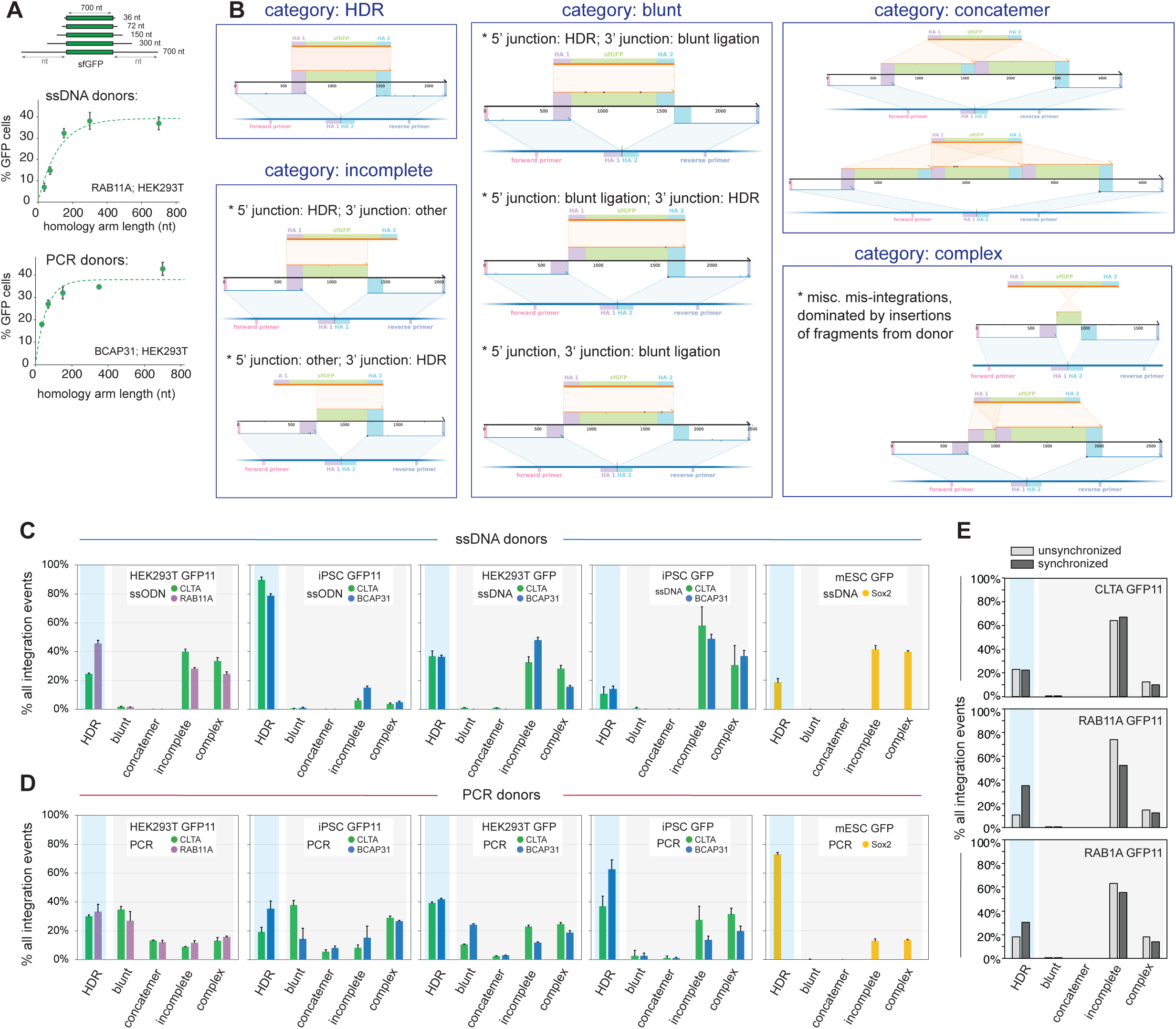
phenotypic classification of repair outcomes. **A.** Titration of homology arm length required for the integration of sfGFP payload in HEK293T cells using ssDNA (top) and PCR donors (bottom). Shown are mean + standard deviation from 3 independent replicate experiments (some error bars are too small to be visible). **B.** Representative examples of read architecture across the 5 different categories of repaired alleles: HDR, blunt, concatemer, incomplete and complex. See text for details. **C-D.** Distribution of integration events across repair categories from cells edited with ssDNA donors (C) or PCR donors (D). Cell-type (HEK293T, iPSC, mESC), payload size (GFP11, sfGFP) and loci targeted (color) are specified. Shown are mean + standard deviation from 3 independent replicate experiments (some error bars are too small to be visible). **E.** Distribution of integration events across repair categories from HEK293T cells with or without cell-cycle synchronization by nocodazole treatment (GFP11 integration with ssODN donors at three separate loci: CLTA, RAB11A and RAB1A).

**Supplementary Figure 3 (related to Fig. 3):**
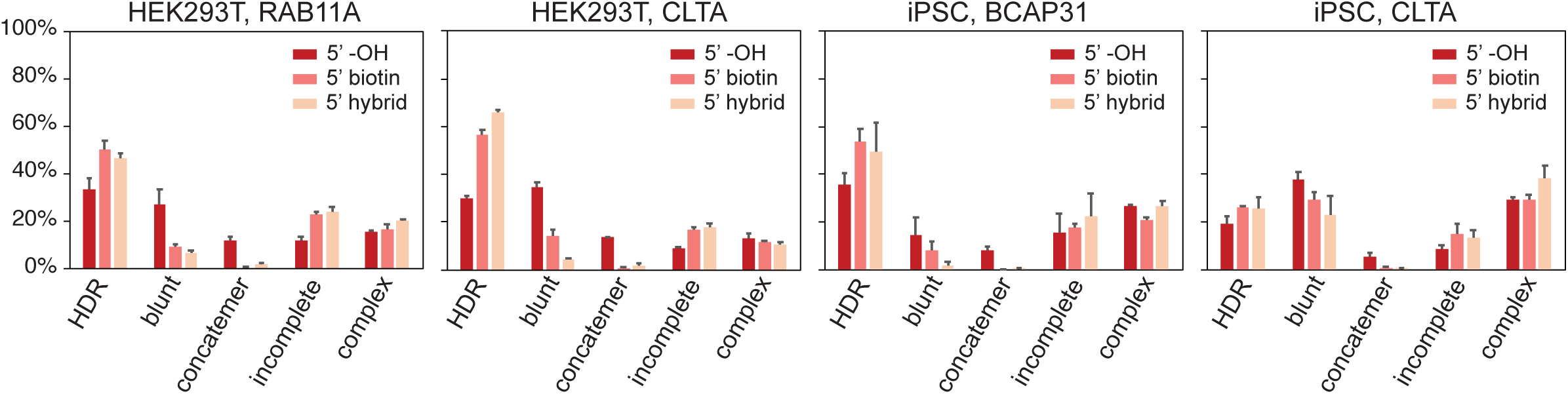
5’ modifications mitigate mis-integration from PCR donors. Distribution of integration events across repair categories from cells edited with GFP11 PCR donors with or without 5’ modifications. Shown are mean + standard deviation from 3 independent replicate experiments (some error bars are too small to be visible).

**Supplementary Figure 4 (related to Fig. 4):**
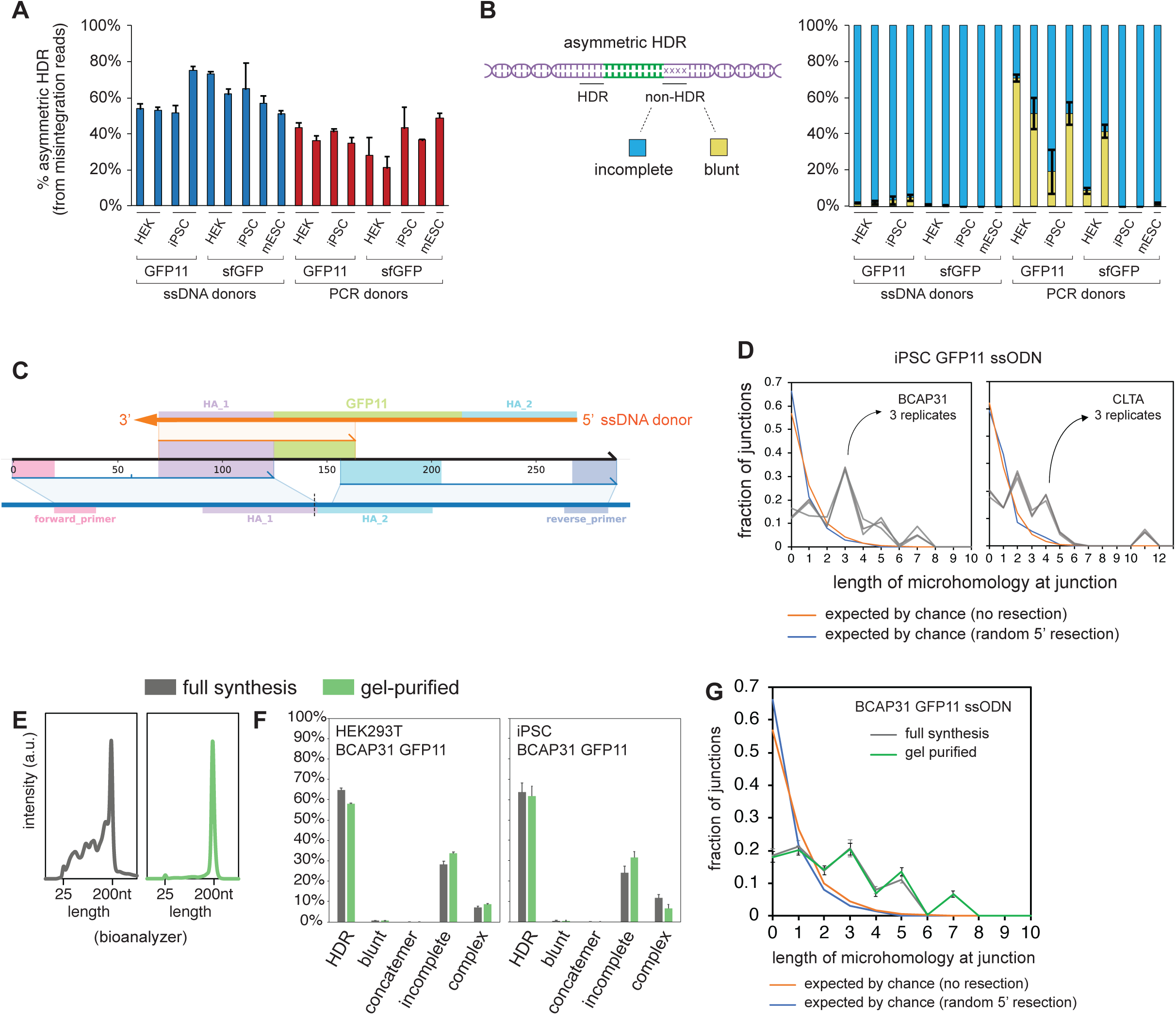
asymmetric HDR. **A**. Prevalence of asymmetric HDR (defined as repair events showing a proper HDR junction at only one homology arm) in mis-integration reads across experimental conditions. Shown are average + standard deviation from 3 independent replicate experiments. Related to Fig 4A. **B.** Identity of non-HDR junction from asymmetric reads: fraction of incomplete vs blunt junctions. Error bars: standard deviation from 3 independent replicate experiments. **C.** Example of knock-knock profile from incomplete read showing microhomology at non-HDR junction. **D.** Related to Fig. 4E. Distribution of microhomology length at incomplete non-HDR junctions across 3 replicate experiments in iPSC (BCAP31 and CLTA loci, MiSeq sequencing). The distributions of microhomology expected by chance considering non-resected or randomly resected DSBs are shown in orange and blue, respectively. **E-G.** Comparison of gel-purified vs. control (full synthesis yield) ssODN donors. Bioanalyzer capillary electrophoresis (**E**) shows that control ssODN include a collection of truncated synthetic products, which can be separated out by gel purification. However, the presence of truncated synthetic products in ssODNs preparations does not explain the complexity of integration outcomes seen in HEK293T or iPSC, as both full synthesis and gel-purified products give rise to very similar integration outcomes (**F**, showing mean + standard deviation from 3 independent replicate experiments). Similarly, the pattern of microhomology at non-HDR junctions

**Supplementary Figure 5:**
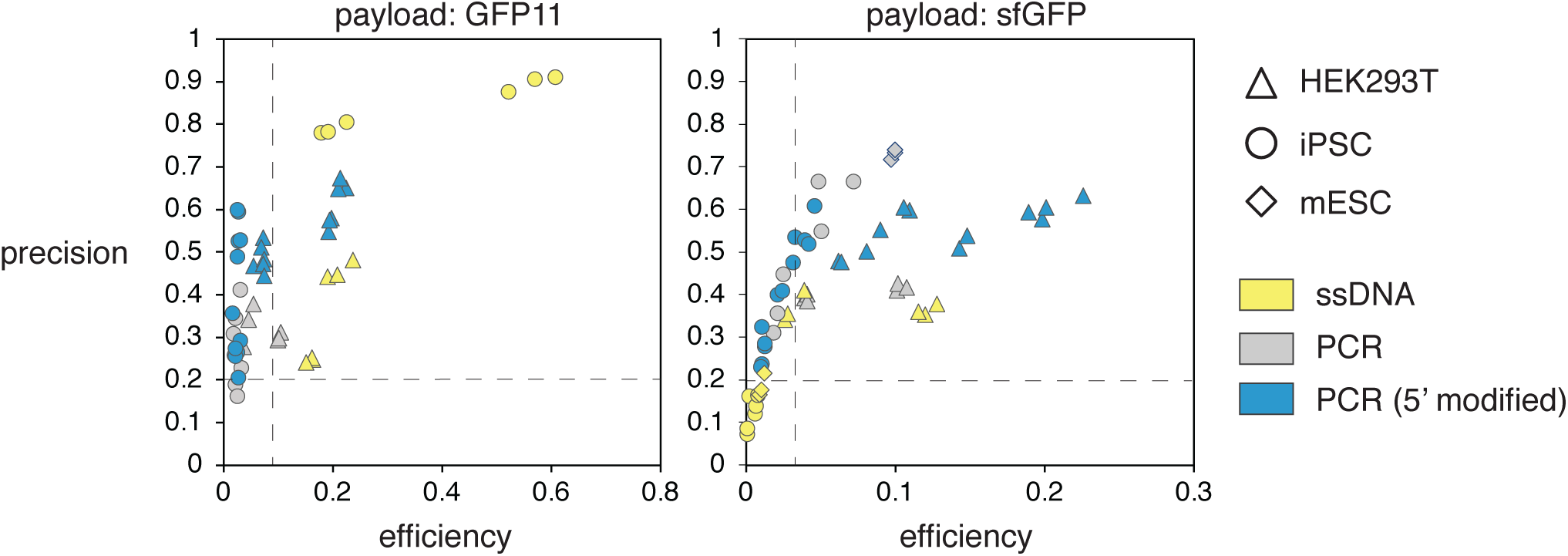
precision vs efficiency of knock-in integration. These graphs represent a summary of all experimental results from our analysis, broken down by cell-types and donor-types. Precision is defined as the ratio of perfect HDR against all payload integration events. Efficiency is defined as the ratio of perfect HDR against all edits (ie. non-wt alleles). Data are separated by payload size (left: GFP11 – small payload, right: sfGFP – large payload). Dashed lines for 20% precision and the fifteenth percentile of maximal efficiency are shown as references.

## Materials and Methods

From “Deep profiling reveals substantial heterogeneity of integration outcomes in CRISPR knock-in experiments” Canaj, Hussmann et al. 2019

### Construct sequences

Sequences for all synthetic constructs, primers, HDR templates and gRNA spacers used are found in Suppl. table 1. Full HDR templates sequences are also provided in Genebank (annotated) and FASTA (unannotated) formats in Suppl. Folder 1.

### Cell culture

#### HEK293T

HEK293T cells were cultured in high-glucose DMEM supplemented with 10% FBS, 1 mM glutamine and 100 µg/mL penicillin/streptomycin (Gibco). HEK293T cells constitutively expressing a GFP1-10 construct were prepared as in ^1^. All cell lines were mycoplasma-free and routinely tested.

#### iPSC

We used the well-characterized human WTC11 iPSC line for all experiments (A gift from Dr. Bruce Conklin, Gladstones Institute; also available through Coriell #GM25256). Cells were maintained on culture plates coated with Matrigel matrix (Corning) and grown in STEMflex media (Gibco). For passaging, cells were detached using StemPro Accutase (ThermoFisher Scientific) and quenched with complete STEMflex media containing 10-µM RHO/ROCK pathway inhibitor (Ri) Y-276932 (STEMCELL Technologies Cat#72304). Cells were subsequently spun down at 300 rcf for 5 minutes and resuspended in media supplemented with Ri, then counted on a cytoFLEX Flow Cytometer (Beckman Coulter). 160,000 cells were seeded in a T25 flask for a 3-day passage or 80,000 cells for a 4-day passage. Media was changed to fresh media without Ri 24 hours after passaging. All cell lines were mycoplasma-free and routinely tested.

To generate a WTC11 line constitutively expressing GFP1-10, TALEN-based knock-in of a CAG promoter-GFP1-10 expression cassette in the AAVS1 safe harbor locus was performed as described in ^2^. A clone was isolated, genotyped for correct insertion at the AAVS1 locus and cytogenic analysis by G-banding was performed to verify its normal karyotype (Cell line Genetics).

#### mESC

The mESC line LF2 and its derivatives were routinely passaged by standard methods in KO-DMEM, 10% FBS, 2 mM glutamine, 1X non-essential amino acids, 0.1mM b-mercaptoethanol and recombinant leukemia inhibitory factor.

### RNP electroporation

#### HEK293T

Cas9/sgRNA RNP complexes were prepared following methods by Lin, Staahl et al. ^3^ with some modifications. S. pyogenes Cas9 protein (pMJ915 construct, containing two nuclear localization sequences) was expressed in E. coli and purified by the UC Berkeley Macrolab following protocols described by Jinek et al. ^4^. HEK293T cells were routinely treated with 200ng/mL nocodazole (Sigma) for 15-18 hours before electroporation to increase editing efficiency as in ^3^. RNP complexes were assembled with 50 pmol Cas9 protein and 65 pmol gRNA just prior to electroporation, and combined with HDR template in a final volume of 10 µL. First, 0.5 µL gRNA (130 µM stock) was added to 2.35 µL high-salt RNP buffer {580 mM KCl, 40 mM Tris-HCl pH 7.5, 20% v/v glycerol, 2 mM TCEP-HCl pH 7.5, 2 mM MgCl2, RNAse-free} and incubated at 70°C for 5 min. 1.25 µL of Cas9 protein (40 µM stock in Cas9 buffer, ie. 50 pmol) was then added and RNP assembly carried out at 37°C for 10 min. Finally, HDR templates and sterile RNAse-free H2O were added to 10 µL final volume. Electroporation was carried out in Amaxa 96-well shuttle Nucleofector device (Lonza) using SF solution (Lonza) following the manufacturer’s instructions. Cells were washed with PBS and resuspended to 10,000 cells/µL in SF solution (+ supplement) immediately prior to electroporation. For each sample, 20 µL of cells (ie. 200,000 cells) were added to the 10 µL RNP/template mixture. Cells were immediately electroporated using the CM130 program, after which 100µL of pre-warmed media was added to each well of the electroporation plate to facilitate the transfer of 25,000 cells to a new 96-well culture plate containing 150µL of pre-warmed media. Electroporated cells were cultured for >5 days prior to analysis.

#### iPSC

Cas9/RNP complexes were assembled with 25 pmol Cas9 protein (EnGen Cas9 NLS, New England Biolabs) and 50 pmol gRNA just prior to electroporation, and combined with HDR template in a final volume of 10 µL. First, 0.5 µL gRNA (100 µM stock) was added to 1.5 µL high-salt RNP buffer {580 mM KCl, 40 mM Tris-HCl pH 7.4, 2 mM DTT, 2 mM MgCl2, RNAse-free} and incubated at 70°C for 5 min. 1.25 µL of Cas9 protein (20 µM stock in Cas9 buffer, ie. 25 pmol) was then added and RNP assembly carried out at 37°C for 10 min. Finally, complexed RNP was further diluted with 1.9 µL of a glycerol-containing buffer {380 mM KCl, 20 mM Tris-HCl pH 7.4, 19.4% v/v glycerol, 1.3mM DTT, 0.9 mM MgCl2, RNAse-free} and HDR templates and sterile RNAse-free H2O were added to 10 µL final volume.

WTC11 iPSC were cultured to 80% confluency. Before electroporation, media was changed to STEMflex media containing 10µM Ri for 2-3 hours. Cells were harvested using accutase, quenched with media containing 10µM Ri and triturated to single cell resolution by pipetting. Singularized cells were harvested, spun down at 300 rcf for 5 minutes, resuspended in STEMflex media containing 10uM Ri to remove accutase, washed in 1X DPBS (Gibco) and resuspended to 10,000 cells/µL in P3 Primary Cell Nucleofector Solution (+ supplement) following manufacturer’s instruction. 20µL of cell resuspension was added to 10µL RNP/donor mixtures prepared as above, and cells were electroporated in an Amaxa 96-well shuttle Nucleofector device (Lonza) using pulse code DS-138. 100µL of pre-warmed media was added to each well of the electroporation plate to facilitate the transfer of 25,000 cells to a new 96-well Matrigel-coated culture plate containing 200µL of pre-warmed media supplemented with 10µM Ri. Media was exchanged to STEMflex alone after 24h. Electroporated cells were cultured for >4 days prior to analysis.

#### mESC

LF2 cells were cultured to 80% confluency and harvested by trypsinization. RNP electroporation was carried out using a Neon system (ThermoFisher). Trypsinized cells were quenched in complete media, washed in 1x DPBS, and resuspended at 10,000 cells/µL in buffer R. 10µL cell resupension was added to 3µL RNP/donor mixtures prepared as above, and electroporation was performed in 10µL Neon tips using the following settings: 1500V, 10ms, 3 pulses. The entire cell sample (100,000 cells) was then transferred to a 24-well plate containing 600µL of pre-warmed media. Electroporated cells were cultured for >5 days prior to analysis.

### Flow cytometry and analysis

Analytical flow cytometry was carried out on a cytoFLEX flow cytometer by Beckman Coulter Life Sciences and cell sorting on a Sony SH800S Cell Sorter. Flow cytometry data analysis and figure preparation was done using the FlowJo software (FlowJo LLC).

### gRNAs

gRNAs were purchased from Integrated DNA technologies (IDT DNA, Alt-R crRNA + tracrRNA).

### Nucleic acid purification with magnetic beads

Magnetic beads for nucleic acid purification were prepared as in ^5^. For a 50 mL working solution of beads, 1mL carboxylate-modified magnetic bead solution (GE Healthcare #65152105050250) is first washed with 3×1mL RNAse-free TE buffer (10mM Tris, 1mM EDTA, pH 8.0) on a magnetic stand. The bead pellet is then resuspended in 50mL DNA precipitation buffer (RNAse-free): {1M NaCl; 10mM Tris-HCl pH 8.0; 1mM EDTA; 18% w/v PEG8000 (Sigma BioUltra); 0.05% v/v Tween20 (Sigma)}. For nucleic acid precipitation and purification, magnetic beads in precipitation solution are added to nucleic acid samples in appropriate ratios and incubated for 10 min at RT. Bead-bound nucleic acids are then isolated on a magnet and washed 2x in RNAse-free 70% EtOH. Beads are air-dried (5-10 min at RT) and the nucleic acids eluted in RNAse-free H2O.

For applications where very large amounts of nucleic acid have to be purified (for example: after in-vitro transcription of long RNA sequences), the stock concentration of magnetic resin in precipitation solution can be increased 3-fold (ie. use 3mL of carboxylate-modified magnetic bead solution for 50mL final working stock).

### HDR donor preparation

#### Short ssODN donors

Short ssODN GFP11 donors templates (Ultramer, <200nt) were purchased from IDT DNA.

#### Long ssDNA donors

ssDNA synthesis was carried out using reverse-transcription (RT) of an RNA intermediate. Thermo-stable TGIRT group II intron reverse transcriptase was chosen due to its superior processivity compared to viral RT enzymes^6^.

ssDNA generation: step 1 – PCR. All constructs used here are amplified by PCR of sequence-verified plasmids. A 100 µL PCR reaction is set using Kapa HotStart HiFi reagents (Kapa Biosystems #KK2601) containing 150 pmol each primer and 10 ng plasmid template. PCR reaction is amplified in a thermocycler: 95°C for 3 min, 30 cycles of {98°C for 20 s, 69°C for 15 s, 72°C for 15 s/kb}, 72°C for 5 min, 4°C final. 2 µL DpnI (NEB # R0176L) is then added and the reaction incubated 1h at 37°C. For purification, 80 µL magnetic beads in precipitation solution are added to the reaction. After magnetic purification, PCR products are eluted in 15 µL of RNAse-free {2 mM Tris-HCl pH 8.0}.

ssDNA generation: step 2 – IVT. A 100 µL IVT reaction is set using HiScribe T7 polymerase (NEB #E2040S). The reaction contained: 10 µL 10x T7 buffer, 2 µL {0.1 M DTT}, 50 µL NTP mix (25 mM each, NEB #N0466S), 2 µL RNAse.In (Promega #N2115), 8 pmol of DNA from step 1 and 10 µL T7 HiScribe enzyme, RNAse-free H2O to 100 µL. The reaction is incubated 3 hours at 37°C, after which 4 µL TurboDNAse (Thermo #AM2238) is added to the reaction and incubated 15 min at 37°C. For purification, 120 µL magnetic beads in precipitation solution are added to the reaction (1.2:1 beads:RNA v:v). We use a 3-fold concentrated stock of magnetic beads in precipitation solution (see above) given the typically very large yield of this reaction. After magnetic purification, RNA products are eluted in 50-100 µL of RNAse-free H2O.

ssDNA generation: step 3 – RT. The RT reaction is set up as follows. First, 400 pmol RNA is mixed with 8 µL of gene-specific RT primer (100 µM in H2O), 15 µL dNTP mix (25 mM each, Thermo #1122) and RNAse-free H2O to 70 µL. To allow for primer annealing, the reaction is incubated 5 min at 65°C and placed immediately on ice for another 5 min. Then are added: 20 µL 5x RNAse-free RT buffer {250 mM Tris-HCl pH 8.3, 375 mM KCl, 15 mM MgCl2}, 5 µL {0.1 M DTT}, 2 µL RNAse.In (Promega #N2115) and 7 µL TGIRTIII enzyme (InGex). The reaction was incubated 1.5 hours at 58°C, after which RNA is hydrolyzed by the addition of 42 µL {0.5 M NaOH, 0.25 M EDTA pH 8.0} and incubated 10 min at 95°C. NaOH is quenched with 42 µL {0.5 M HCL}. For purification, 0.9X v:v magnetic beads in precipitation solution are added to the reaction. After magnetic purification, ssDNA products are eluted in 25 µL sterile H2O. The reaction can be scaled down two-fold to fit in PCR strip format. Typical yields: 50-200 pmol ssDNA per 500 pmol RNA template.

#### PCR dsDNA donors

All PCR donors include homology templates (payload + homology arms) flanked by universal handle sequences used for PCR amplification (left handle: 5’-GGGAACCTCTTCTGTAACTCCTTAGC-3’; right handle: 5’- CCTGAGGGCAAACAAGTGAGCAGG -3’). Donors were amplified from corresponding plasmid templates using these universal handles as primers. PCR primers were ordered from IDT DNA either unmodified (“5’ OH”) or with the following 5’ modifications (exact primer sequences are found in Supplementary table 1):

“5’ biotin”: biotin + 5x phosphoro-thioate bonds to increase in vivo stability.

“5’ hybrid”: artificial nucleotide sequence (to serve as a ssDNA overhang) + 6xPEG linker.

#### Plasmid donors

Plasmid donors were prepared using Zymopure Maxipreps kits (Zymo Research) and further purified for endotoxin removal through Endopure spin columns (Zymo Research), following manufacturer’s instructions.

### Amplicon sequencing

#### gDNA preparation

gDNA was extracted by cell lysis using QuickExtract DNA Extraction Solution (Lucigen). From a confluent culture in 96-well plate, media was removed, cells were washed 1x in DPBS and resuspended in 50 µL QuickExtract. The cell layer was detached by repeated pipetting and transferred to a PCR plate for incubation. The lysate was incubated as follows {65°C for 20 min, 98°C for 5min, 4°C final}. gDNA was used directly from this preparation.

#### Illumina Miseq amplicon sequencing

Amplicon Libraries were created using a two-step PCR protocol: the first PCR amplifies the target genomic locus and adds universal amplification handle sequences, while the second PCR introduces index barcodes using the universal handles.

PCR1: this PCR uses a “reverse touchdown” method designed to accommodate a number of different annealing temperatures for a number of different targets. 50-µL PCR reactions were set using 2x KAPA HiFi Hotstart reagents (Roche) with 2µL extracted gDNA, 80pmol each primer and betaine to 0.8M final concentration. PCR conditions: 95°C 3min; 3 cycles of {98°C for 20s, 63°C for 15s, 72°C for 20s}, 3 cycles of {98°C for 20s, 65°C for 15s, 72°C for 20s}, 3 cycles of {98°C for 20s, 67°C for 15s, 72°C for 20s}; 17 cycles {98°C for 20 s, 69°C for 15 s, 72°C for 20s} then 72°C for 1min; 4°C final.

PCR2: amplicons were diluted 1:100 and 1 µL was used into a 40-µL barcoding reaction using 20 µL 2x KAPA HiFi Hotstart reagents (Roche) and 80pmol each barcoded primer. Thermocyling conditions: 95°C 3min and 12 cycles of {98°C for 20s, 68°C for 15s, 72°C for 12s} then 72°C for 1min; 4°C final.

Barcoded amplicons were analyzed using capillary electrophoresis (Fragment Analyzer, Agilent), pooled and purified using magnetic beads (beads:DNA 0.8:1 v:v). Sequencing was performed on an Illumina Miseq V3 platform using standard P5/P7 primers.

#### Pacific Biosciences Library Preparation

Amplicon libraries were prepared from extracted gDNA by 1-step PCR using barcoded genomic primers. 100-µL PCR reactions were set using 2x KAPA HiFi Hotstart reagents (Roche) with 1µL extracted gDNA, 100pmol each primer and betaine to 1M final concentration. PCR conditions for 3-5kb amplicons: 95°C 3min; 27 cycles {98°C for 20 s, 69°C for 20 s, 72°C for 5:23min} then 72°C for 10min; 4°C final. PCR conditions for 2kb amplicons 95°C 3min; 27 cycles {98°C for 20 s, 67°C for 20 s, 72°C for 2:30min} then 72°C for 5min; 4°C final.

Amplicons were purified using magnetic beads (beads:DNA 0.8:1 v:v), analyzed using capillary electrophoresis (Fragment Analyzer, Agilent) and pooled. Pooled libraries were capped with SMRT adapters using SMRTbell Template Prep Kit 1.0 (Pacific Biosciences) and the Sequel Binding Kit 3.0 was used to bind primers and polymerase before sequencing. Sequencing was performed using the Sequel System using the SMRT Cell 1M v3 and acquiring 20h movies.

#### SMRT consensus sequence analysis

Demultiplexing and circular consensus sequence analysis and FASTQ generation from SMRT polymerase reads was performed using the SMRTlink 6.0 software (Pacific Biosciences). To provide appropriate computing power, we used cloud computing (AWS, Amazon Web Services. Details regarding AWS setup for individual downloading and running SMRTlink 6.0 can be found in this public Gist on GitHub: https://gist.github.com/hcanaj/3a5bb9e8357016181a6a19a840428679

Analysis was subsequently conducted via the the SMRTlink 6.0 GUI interface. Reads were demultiplexed using the RSII_384_barcodes set using the following parameters: {Same Barcodes: TRUE; Minimum barcode score: 45; Infer barcodes: TRUE}. Routine CCS Analysis was performed after demultiplexing using the following parameters:{Minimum number of passes: 3; CCS Accuracy 0.99}.

### Data analysis with knock-knock

Full instructions for usage of knock-knock and full source code are available at https://github.com/jeffhussmann/knock-knock. knock-knock can be installed from bioconda^7^. knock-knock relies on numpy^8^, pandas^9^, matplotib^10^, samtools^11^, pysam (github.com/pysam-developers/pysam), blastn^12^, STAR^13^, minimap2^14^, and GNU parallel^15^.

#### For Illumina data: reconstructing full reads from paired-end runs

Before generating alignments to reference sequences, Illumina data is preprocessed by stitching together each overlapping read pair into a single sequence. This is done by aligning R1 to the reverse complement of R2 after prepending the expected Illumina sequencing primers to each read. Read pairs for which a confident overlapping region can be identified are stitched together, with the higher quality read base retained for any position in the overlapping region at which the two reads disagree. Read pairs for which no confident stitching alignment can be identified (either because the amplicon was longer than the combined length of the read pair, or because sequencing errors introduced too many discrepancies between R1 and R2) are retained and processed separately.

#### Alignment generation

knock-knock generates local alignments between each sequencing read and relevant reference sequences in two stages. First, in order to maximize sensitivity of alignment to the sequences most expected to participate in the repair process, blastn is used to align a read to the targeted genomic sequence in the immediate vicinity of the cut site (by default, from 500nt upstream of the upstream amplicon primer to 500nt downstream of the downstream amplicon primer, referred to hereafter as the target sequence) and to any donor sequence(s) present. (For full blastn parameters, see knock-knock/blast.py.) Next, more computationally efficient tools are used to align to the full targeted genome and to any other specified supplemental genome: STAR for Illumina data, and minimap2 for PacBio data. (For full STAR and minimap2 parameters, see knock-knock/experiment.py.) All alignments produced for a given read by both stages are collected and passed into the categorization process.

#### Categorization

Categorization for a given read begins with a set of local alignments identified by several different alignment strategies. knock-knock examines the relative positioning of these alignments on the read and on the various reference sequences to place the read into one of several categories, each of which can have one or more subcategories. For a complete delineation of the decision tree, see the source code of Layout.categorize() in knock-knock/layout.py.

Alignments are first refined by splitting each alignment into multiple smaller local alignments at any large insertion or deletion or any cluster of high total edit distance in a short window. (Large insertions and deletions are defined to be greater than or equal to 3 nts for Illumina data or greater than or equal to 5 nts for PacBio data. A high edit distance cluster is defined to be more than 5 edits in an 11nt window for Illumina data; high edit distance windows are not split at for Pacbio data). Alignments are further refined by making allowances for irregularities that may be present in primer sequences as a result of imperfections in primer synthesis or in priming specificity. First, because oligonucleotide primers frequently contain 1nt deletions, initial alignments to primer regions may be incorrectly truncated if there aren’t enough matching bases after the deletion to offset the scoring penalty of the deletion. To correct these truncated alignments, if any alignment ends close to but not at a read end, it is repeatedly extended by including a 1nt deletion and any subsequent matches between the read and reference past this deletion until reaching a potential deletion that would not expose any additional matches. Second, if the set of alignments produced by this process does not cover either end of the sequencing read, each uncovered edge is re-aligned to the expected amplicon primers using a custom Smith-Waterman implementation. This allows detection of chimeric sequences produced by off-target amplification of unintended regions of the targeted genomic by the amplicon primers.

After refining alignments, a read is checked for primer-containing edge alignments: alignments to the targeted sequence that include one of the amplicon-primer-binding regions and extend to within 10nt of one end of the read. (Note that a single alignment can be a primer-containing edge alignment for both primers if it covers the entire read). If there exist primer-containing edge alignments for both amplicon primers that are in the same orientation and cover opposite ends of the read, the read is considered to have a well-formed layout. If not, it is categorized as ‘malformed layout’. Subcategories of ‘malformed layout’ are ‘too short’, consisting of reads less than 50nt long that are typically interpreted as primer-dimers; ‘no alignments detected’; ‘extra copy of primer’, consisting of read for which there are multiple disjoint alignments to at least one of the amplicon primers, typically as a result of incorrect PacBio CCS formation; ‘missing a primer’, consisting of reads for which at least one of amplicon primers did not have any alignments that contained it; and ‘primer far from read edgè and ‘primers not in same orientation’, consisting of reads for which alignments to primers were not positioned as expected on the read or oriented as expected.

For well-formed layouts, the extent of the read covered by the primer-containing edge alignments is then computed. If the entire read is covered, there was no integration event. For well-formed reads with no integration, if the primer-containing edge alignments are not the same single alignment, the two alignments are joined with a deletion and merged into a single alignment. If the single alignment spanning the two amplicon primers doesn’t contain any indels within 10nt of the expected cut site, the read is categorized as ‘WT’. If a single spanning alignment contains a single insertion or deletion within 10nt of the cut site, the read is categorized as a ‘simple indel’, with subcategories ‘insertion’, ‘deletion <50nt’, and ‘deletion >=50nt’. If a single spanning alignment contains multiple indels within 10nt of the cut site, the read is categorized as ‘uncategorized’ with subcategory ‘multiple indels near cut’. If a single spanning alignment contains a mismatch between the read and target genome with 10nt of the cut, the read is categorized as ‘uncategorized’ with subcategory ‘mismatch(es) near cut’.

If primer-containing edge alignments do not cover the whole read, there is integrated sequence to be accounted for. The portion of the read not covered by the primer-containing edge alignments is called the integration gap. If a distinct alignment to the target sequence within 500nt of the amplicon primers covers the integration gap, the read is categorized as ‘complex indel’ with subcategory ‘templated insertion’. If the integration gap is mostly covered by a single alignment to a homologous donor sequence, with at most 10nt total of gap uncovered and with no large indels in the donor alignments, the gap is considered to be covered by the donor and the junctions between the primer-containing edge alignments and the donor alignment are examined to further classify the integration (see below). If multiple disjoint alignments to the donor collectively cover the integration gap, these alignments are examined to see if they consist of concatamerizations of the donor sequence. If all junctions between donor alignments in the integration gap extend to within 1nt of the edge of the donor sequence and leave at most 2nt of unexplained read sequence between the donor alignments, the integration is flagged as a donor concatemer and sent to target junction classification (see below). If neither of these conditions is met and there exists any alignment to a non-homologous donor sequence that overlaps the integration gap, the read is categorized as ‘non-homologous donor’. If a single non-homologous donor alignment leaves at most 2nt of the integration gap unexplained, the read has subcategory ‘simple’, otherwise it has subcategory ‘complex’. If neither primer-containing edge alignments extends at least 20nt past the primer and the entire integration gap is covered by a single alignment to the targeted genome that is not within 500nt of the amplicon primers, the read is interpreted as being produced by nonspecific binding of the amplicon primers to unintended alternative locations in the target genome, and the read is categorized as ‘malformed layout’ with subcategory ‘nonspecific amplification’. If at least one primer-containing edge alignment does extend substantially past the primers and the integration gap is covered by a single alignment to the target genome far from the amplicon primers, the read is interpreted as integration of distal genomic sequence, and the read is categorized as ‘genomic insertion’.

For reads with integration flagged as from a single or concatemerized homologous donor, the junctions between the donor alignment(s) and the on each side are separately analyzed. If the target sequence cleanly transitions to donor sequence at a junction, with one full length copy of the relevant homology arm present in the read and with no indels greater than or equal to 3 nts long within the first nts of the donor sequence past the homology arm, the junction is classified as a clean handoff. If the junction is not clean and the donor alignment extends all the way to within 1 nt of an edge of the donor sequence, the junction is classified as blunt. Otherwise, the junction is classified as imperfect. A read with a single integration gap donor alignment with clean handoffs at both junctions is categorized as ‘HDR’; with at least one blunt junction is categorized as ‘blunt mis-integration’, with a subcategory that records which of the two junctions is blunt; with one junction that is a clean handoff and the other that is imperfect as ‘incomplete HDR’, with a subcategory that records which junction is clean and which is imperfect; and with any other combination of junctions as ‘complex mis-integration’. A read with a concatemer integration is classified as ‘concatenated mis-integration’ with a subcategory that describes the two junctions, unless the donor is a plasmid, in which case concatamerization indicates alignments that collectively cross the arbitrary point at which the circular plasmid sequence was linearized to create the donor reference sequence. In this case, the read is categorized as ‘incomplete HDR’.

Reads that do not satisfy any of these conditions but have any alignment to a donor sequence that overlaps the integration gap are categorized as ‘complex mis-integration’. All other reads are categorized as ‘uncategorized’.

#### Visualization

A full tour of the visualizations produced by knock-knock can be found at https://github.com/jeffhussmann/knock-knock/blob/master/docs/visualization.md. Diagrams of local alignments are generated using matplotlib. See ReadDiagram in knock-knock/visualize.py for complete source code. Interactive outcome tables and interactive outcome-specific length distribution plots are generated using matplotlib, pandas, jupyter (doi:10.3233/978-1-61499-649-1-87), and Bootstrap (github.com/twbs/bootstrap).

### Quantification of microhomology at repair junctions

For reads categorized as incomplete HDR, the length of microhomology at each junction between an alignment to the donor and primer-containing edge alignment to the target is calculated by identifying the segment of the read (if any) covered by both the target alignment and donor alignment. To prevent the potential incorrect inference of microhomology at overlaps produced by incorrect extension of alignments past the true end of donor or target sequence, the initially identified overlap is trimmed back to the first indel or mismatch that occurs from 5nt before the start of the overlap onward. The length of overlap remaining after applying this trimming process from both sides is taken to be the microhomology length.

To calculate the expected distribution of microhomology lengths that would be observed at random junctions between target and donor for a given combination of target cut site and donor sequence, two separate cases are treated: one in which the target sequence extends all the way to cut site while the donor sequence is randomly truncated, and one in which both the target sequence and donor sequence are randomly truncated. For each configuration, for each of the two homology arms, the sequence of the target on the side of the cut containing that homology arm, offset by a random length (either 0 for the non-truncated case, or every integer between 0 and 100 for the resected case), is lined up against the sequence of the donor, offset from the edge on the side containing the homology arm by a every integer between 0 and the length of the donor sequence), and the total number of matching nucleotides surrounding the resulting junction between target and donor is counted. The distribution of lengths for all such offset combinations are recorded, excluding offset combinations that result in alignment of the intended homology arms with each other.

## Notes

https://github.com/jeffhussmann/knock-knock

